# IP_3_-mediated Ca^2+^ transfer from ER to mitochondria stimulates ATP synthesis in primary hippocampal neurons

**DOI:** 10.1101/2025.06.28.661766

**Authors:** Ankit Dhoundiyal, Vanessa Goeschl, Stefan Boehm, Helmut Kubista, Matej Hotka

## Abstract

During electrical activity, Ca^2+^ enhances mitochondrial ATP production, helping to replenish the energy consumed during this process. Most Ca^2+^ enters the cell via ligand– or voltage-gated channels on the neuronal membrane, where it stimulates the release of additional Ca^2+^ from the endoplasmic reticulum (ER). Although the influence of cytosolic Ca^2+^ on neuronal metabolism has been widely investigated, relatively few studies have explored the contribution of ER Ca^2+^ release in this context. Therefore, we investigated how activity-driven Ca^2+^ crosstalk between the ER and mitochondria influences the regulation of mitochondrial ATP production. We show that in primary hippocampal neurons derived from rat pups of either sex, depletion of ER Ca^2+^ led to a reduction in mitochondrial Ca^2+^ levels during both resting and stimulated states, while exerting only a minimal impact on cytosolic Ca^2+^ levels. Additionally, impaired ER-mitochondria Ca²⁺ transfer led to a reduction in mitochondrial ATP production. Similar effects were observed when inositol-3-phosphate receptors (IP_3_Rs), but not ryanodine receptors (RyRs), were pharmacologically inhibited. Together, our findings show that, in hippocampal neurons, Ca^2+^ is transferred from the ER to mitochondria through IP_3_ receptors, and this Ca^2+^ crosstalk in turn enhances mitochondrial ATP production in response to neuronal activity.

**Highlights:**

1. Ca^2+^ adjusts mitochondrial ATP synthesis to neuronal activity
2. In the neuronal somata ER-mitochondria Ca^2+^ crosstalk occurs via IP_3_ receptors
3. IP_3_-mediated Ca^2+^ release occurs across a wide range of firing intensities.

## 1. Introduction

Neuronal electrical activity is an energy-demanding process. The energy stored as ATP is utilized to sustain synaptic neurotransmission by preserving ionic gradients across the membrane, replenishing and recycling synaptic vesicles, and reutilizing released neurotransmitters by re-uptake. (Harris et al., 2012; Li & Sheng, 2022; Rangaraju et al., 2014). The ATP demand created by these processes is preferentially met by mitochondria which boost their ATP production in response to increases in cytosolic Ca^2+^ caused by neuronal activity (Contreras & Satrústegui, 2009; del Arco et al., 2023; Denton et al., 1980; Llorente-Folch et al., 2013).

Extracellular Ca^2+^ enters the cytosol predominantly via ligand-gated and voltage-gated Ca^2+^ channels (VGCCs) (Berridge, 1998; Brini et al., 2014), out of which L-type Ca^2+^ channels (LTCCs) are gaining increasing attention (Berger & Bartsch, 2014; Catterall, 2011; Hotka et al., 2020). LTCCs are expressed in neurons in the somato-dendritic compartment (di Biase et al., 2011; Stanika et al., 2016), where they mediate Ca^2+^ influx in response to excitatory postsynaptic potentials (Mermelstein et al., 2000; Striessnig et al., 2014). Owing to their ability to transmit synaptic Ca^2+^ signals along the dendrites to neuronal cell bodies, LTCCs have been shown to stimulate somatically located mitochondria in multiple neuronal subtypes (Hotka et al., 2020; Zampese et al., 2022). However, details of the coupling between LTCCs and mitochondria are not fully understood.

The endoplasmic reticulum (ER), an integral Ca^2+^ store spanning throughout a neuron, is posed ideally to mediate this interaction. Upon excitation, Ca^2+^ is released from the ER in a process called calcium-induced calcium release (CICR) (Berridge, 1998; Heine et al., 2020). Mitochondria form functional complexes with the Ca^2+^ release channels of the ER in regions of contact between the two organelles known as mitochondria-associated membranes (MAMs) (Rizzuto et al., 1998; Wu et al., 2017).

While the importance of Ca^2+^ transfer from ER to mitochondria in promoting mitochondrial ATP synthesis has been demonstrated in several non-neuronal cell types (Bartok et al., 2019; Cárdenas et al., 2010, 2016; Díaz-Vegas et al., 2018; Filadi et al., 2018), only a few studies have addressed this mechanism in neurons, and the results have remained inconsistent (Pérez-Liébana et al., 2022; Zampese et al., 2022; Liiv et al., 2024.).

One challenge in the analysis of a role of ER Ca^2+^ release in the control of mitochondrial bioenergetics is the fact that Ca^2+^ can stimulates mitochondria at various sites accessible from inside and outside of the organelle (del Arco et al., 2023; Díaz-García et al., 2021; Griffiths & Rutter, 2009; Llorente-Folch et al., 2013, 2015; Satrústegui et al., 2007; Szibor et al., 2020); the relative contribution of these different pathways depends on the intensity of neuronal activity (Dhoundiyal et al., 2022, Groten & MacVicar, 2022; Stoler et al., 2022). Moreover, individual Ca^2+^-sensitive pathways may substitute for each other, thereby providing neurons with substantial metabolic flexibility. Accordingly, inhibition of a single Ca^2+^-sensitive pathway involved in the regulation of the metabolic machinery in neurons may fail to impact mitochondrial ATP synthesis (Dhoundiyal et al., 2022). Likewise, deletion of the main mitochondrial Ca^2+^-entry pathway, the mitochondrial calcium uniporter (MCU), in mice did not lead to a strong metabolic phenotype (Luongo et al., 2015; Pan et al., 2013; P. Wang et al., 2020). Under this condition, redox shuttles sensitive to fluctuations of cytosolic Ca^2+^ can compensate for the loss of MCU (Zampese et al., 2022). Thus, the redundancy of multiple mechanisms by which Ca^2+^ can boost oxidative metabolism in mitochondria complicates the interpretation of experimental results.

In this study, we sought to monitor the ER-mitochondria Ca²⁺transfer in response to neuronal electrical activity directly, and to determine how individual ER Ca²⁺-release channels may contribute to the stimulation of mitochondrial ATP synthesis. The ER–mitochondria Ca^2+^ crosstalk was studied by monitoring cytosolic calcium ([Ca^2+^]_i_), ER Ca^2+^, mitochondrial Ca^2+^ ([Ca^2+^]_mito_), and mitochondrial ATP synthesis in individual hippocampal neurons. To eliminate the potentially confounding influence of Ca^2+^-sensitive redox shuttles, key experiments were performed under conditions where neurons depended exclusively on mitochondrial calcium uptake to enhance their ATP production. All parameters were monitored in both spontaneously firing neurons and during neuronal activation induced by electric field stimulation (EFS). Our results demonstrate that calcium release from the ER triggers an increase in mitochondrial calcium through an IP_3_-mediated pathway, which in turn promotes mitochondrial ATP production. This ER–mitochondria calcium crosstalk serves to adapt mitochondrial ATP synthesis to the metabolic burden created by neuronal activity.

## 2. Materials and Methods

### 2.1 Drugs and Chemicals

Table 1 lists the pharmacological agents used, together with their final concentrations and catalogue numbers. DMSO was added to the control solutions as needed to match its concentration with that of the experimental solutions. In several tests, cells were pretreated for either one hour (2DG + Pyruvate, Ryanodine, and CPA) or for 15 minutes (2-APB, U73122, and U73343). In experiments involving local perfusion of the cells, the compounds were added to the superfusion solutions.

**Table 1:** Compound list.

| Compound Name | Final Concentration | Catalog number | Company |
| --- | --- | --- | --- |
| 2-APB | 30 $\mu\text{M}$ | catalog#100065 | Sigma |
| 2-Deoxy-D-glucose (2DG) | 2 mM | catalog #D8375 | Sigma |
| Calcimycin | 20 $\mu\text{M}$ | catalog#CAY11016 | Cayman Chemical |
| CPA | 100 $\mu\text{M}$ | catalog #C1530 | Sigma |
| Dimethylsulfoxide (DMSO) | 0.01% | catalog #D2650 | Sigma |
| EGTA-AM | 1 mM | catalog #CAY-20401-10 | Cayman Chemical |
| Ionomycin | 20 $\mu\text{M}$ | catalog #I0634 | Sigma |
| Isradipine | 1 $\mu\text{M}$ | catalog #I6658 | Sigma |
| Ryanodine | 50 μM | catalog #1329 | Tocris |
| Sodium pyruvate | 2 mM | catalog #P8574 | Sigma |
| U73122 | 1 μM | catalog#U6756 | Sigma |
| U73343 | 1 μM | catalog#4133 | Tocris |

### 2.2 Hippocampal neuron primary cell culture

Pregnant Sprague-Dawley rats were obtained from Charles River Laboratories (Sulzfeld, Germany). Neonatal animals of either sex were killed by decapitation in full accordance with all rules of the Austrian animal protection law for details see (Hotka et al., 2020). Brains were removed to dissect the hippocampi in ice-cold buffer. Primary co-cultures of hippocampal neurons and glial cells were prepared after enzymatic digestion of the tissue with papain and mechanical dissociation with Pasteur pipettes (trituration) as described previously (Kubista et al., 2025). Neurons were cultured for at least 14 days at 37°C and 5% CO2 in Dulbecco’s modified Eagle’s medium – high glucose (20 mM glucose) (D5796, Sigma-Aldrich) supplemented with 10% γ-irradiated fetal bovine serum (S 0415, Biochrom).

### 2.3 Transfections

Transfections were conducted using Lipofectamine 2000 reagent (Thermo Fisher Scientific) in accordance with the manufacturer’s instructions, with the following modification: Neuronal cultures older than 14 days in vitro were incubated in 500 µl of antibiotic-free medium, supplemented with 3 µl of Lipofectamine 2000 and 1 µg of plasmid DNA, for a duration of 3 hours. The transfection medium was substituted with the original medium, and cultures were maintained in the incubator for 3 more days. Experiments were conducted on the third day post-transfection.

### 2.4 Viral transductions

All viral transductions were performed using adeno-associated viruses (AAV) in vitro by direct addition of viral particles into culture dishes. The optimal amount of virus was estimated by titration and typically 1 µl of AAV particles was used per 2.7 ml of culture media. Neurons were used for the experiment 9 days after transduction.

### 2.5 Electric field stimulation

Field stimulation was carried out using two platinum electrodes, each with a diameter of 1 mm, positioned 10 mm apart within the culture dishes. An external voltage of 20 V amplitude and 1.5 ms duration was applied using a stimulator (model S44, Grass Medical Instruments), which facilitated the generation of single pulses and trains of pulses at specified frequencies. Measurements were conducted on neurons located centrally between the two electrodes.

### 2.6 Imaging

Cells were cultured in glass-bottom dishes (model P35GC-1.5–14–C, MatTek). Before each experiment, the culture medium was replaced with an external solution composed of (in mM): NaCl (140), KCl (3), CaCl_2_ (2), MgCl_2_ (2), HEPES (10), glucose (20), with NaOH utilized to adjust the pH to 7.4. For drug application, we used the Octaflow II system, which comprises 8 reservoirs and a micro manifold featuring eight channels that converge at a quartz outlet with a diameter of 100 µm. The experiments were performed at room temperature, with continuous superfusion of the cells. In certain instances, neurons were incubated in pharmacological agents as required either before and/or during the experiments. Neurons were imaged using a laser scanning confocal microscope (Nikon, model A1R) equipped with a focus clamp. Fluorescent indicators were employed to assess the metabolic activity of neurons.

ATP/ADP ratio: PercevalHR, a genetically encoded fluorescence biosensor was used to quantify the cytosolic ATP/ADP ratio in cells (Tantama et al., 2013). PercevalHR was introduced into cells using AAV-mediated transduction. Excitation wavelengths of 403 nm and 488 nm were used to excite the fluorophore and the resulting emission was detected at 525 nm. Fluctuations in the cytosolic ATP/ADP ratio were monitored by the F488/F403 ratio. The fluorescent values (F) obtained in each experiment were normalized to the value obtained from the pre-stimulation region (F_0_). All Perceval HR data are represented as averaged normalized data (F/F_0_) traces accompanied by standard error of the mean (SEMs). GW1-PercevalHR was a gift from Gery Yellen, Department of Neurobiology, Harvard Medical School, Boston, USA (plasmid #49082, Addgene).

Cytosolic Ca^2+^[*Ca*^2+^]_i_: Fluo-4 AM (catalogue #F14201, Thermo Fisher Scientific) was used to assess the cytosolic Ca^2+^ concentration. The cells were loaded with 1 μM Fluo4 AM for 15 minutes at 37°C followed by a washout with extracellular solution. An excitation wavelength of 488 nm and a detection wavelength of 525 nm were used for imaging. Following each experiment, the cells were permeabilized with ionophores (20 μM ionomycin+20 μM calcimycin), and the maximum and minimum fluorescence (F_max_ and F_min_) for each cell was ascertained by administering 5 mM Ca^2+^, followed by 0 mM Ca^2+^+1 mM EGTA, respectively (Figure S1). Subsequently, [Ca^2+^]_i_ for different time points was calculated using the formula:

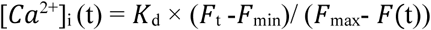

where [*Ca*^2+^]_i_(t) is the cytosolic Ca^2+^ concentration at time t; F(t) is the fluorescence value at time point t; F_min_ is the minimum fluorescence value recorded under 0 mM Ca^2+^+1 mM EGTA; F_max_ is the maximum fluorescence value recorded under 5 mM Ca^2+^; and *K*_d_ is the dissociation constant for Fluo4 AM (K_d_=325 nM).

Mitochondrial Ca^2+^ [*Ca*^2+^]_mito_: The genetically encoded fluorescent indicator CEPIA2mt was employed to monitor mitochondrial Ca^2+^ (Suzuki et al., 2014). The expression vector pCMV CEPIA2mt was a gift from Masamitsu Iino (plasmid #58218, Addgene). CEPIA2mt was expressed in cells using AAV-mediated transduction. Cells were excited with a 488 nm wavelength, and emitted fluorescence was detected at 525 nm. At the end of each experiment, ionophores (20 μM ionomycin+20 μM calcimycin) were used to permeabilize the cells, and the maximum and minimum fluorescence (F_max_ and F_min_) for each cell was ascertained by the addition of 5 mM Ca^2+^, followed by 0 mM Ca^2+^ + 1 mM EGTA (Figure S2). Subsequently, [Ca^2+^]_mito_ at different time points was calculated utilizing the formula:

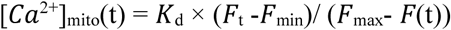

where [*Ca*^2+^]_mito_ (t) is the mitochondrial Ca^2+^ concentration at time t; F is the fluorescence value at time point t; F_min_ is the minimum fluorescence value recorded under 0 mM Ca^2+^+1 mM EGTA; F_max_ is the maximum fluorescence value recorded under 5 mM Ca^2+^; and *K*_d_ is the dissociation constant for CEPIA2mt (K_d_=160 nM).

ER-Ca^2+^: To monitor Ca^2+^ levels in the endoplasmic reticulum, neurons were transfected with the genetically encoded fluorescent sensor CEPIA1er. The expression vector pCMV R-CEPIA1er was a gift from M. Iino (Addgene plasmid no. 58216). Cells were excited with 562 nm wavelengths, and the emitted fluorescence was detected at 595 nm.

PIP_2_: To monitor PIP_2_ levels, we used the pleckstrin homology (PH) domain of phospholipase Cδ (Heo et al., 2006), N-terminally tagged with BFP, which specifically binds to PIP2 in the plasma membrane. A decrease in membrane PIP_2_ is indicated by the translocation of the fluorescence signal from the membrane to the cytosol (Losgott et al., 2025). Cells were excited with 405 nm wavelengths, and the emitted fluorescence was detected at 525 nm.

### 2.7 Data analysis and statistics

Fluorescence intensity was quantified as the average intensity from a specified region of interest located at neuronal somata. The graphs present averaged data traces accompanied by standard error of the mean (SEMs). In the experiments using Fluo4 AM and CEPIA2mt, the fluorescence measurements were transformed into actual Ca^2+^ concentration values (in nM) and subsequently plotted accordingly, whereas in experiments using Perceval HR and CEPIA1_er_ data are presented as normalized values.

For statistical analysis, the Shapiro–Wilk normality test was used to determine whether the data follow a normal distribution. If allowed by the normality tests, parametric tests were used. When comparing values between two conditions, a paired or unpaired t-test was employed, depending on the experimental design. When comparing values between three or more experimental conditions, a standard one-way ANOVA with Tukey’s multiple comparisons was employed. For non-parametric analyses, Mann–Whitney test or Kruskal–Wallis test was used. The statistical analysis was conducted using the 9.1.2 version of GraphPad Prism software. p values smaller than 0.05, 0.01, 0.001 and 0.0001 are indicated by one, two, three and four asterisks, respectively. Differences that are not statistically significant (p values ≥ 0.05) are labelled as ns (not significantly different). n values denote the total number of individual neurons from which data were collected. All of the data used in this investigation came from at least two different independent cell culture preparations. Each preparation contained neurons derived from the hippocampi of two female and two male rat pups.

## 3. Results

### 3.1 EFS-induced rises in cytosolic Ca^2+^ originate primarily from the extra-rather than intracellular compartment

Elevation of cytosolic calcium due to electrical activity in neurons can result from transmembrane Ca^2+^ entry into the cytoplasm which in turn leads to release of Ca^2+^ from intracellular stores such as ER (Berridge, 1998; Berridge et al., 2000). The primary activity-dependent pathway linking extracellular calcium influx to ER-Ca^2+^ release is provided by LTCCs (Verkhratsky & Shmigol, 1996; Vierra et al., 2021) which thereby coordinate various neuronal functions. These include the regulation of activity-dependent gene transcription on one hand and stimulation of neuronal metabolism on the other. For the regulation of gene transcription, LTCCs transmit Ca^2+^ signals to the nucleus in an ER-independent manner (Wild et al., 2019). LTCC-mediated stimulation of mitochondrial ATP synthesis, however, depends on release of Ca^2+^ from the ER (Hotka et al., 2020).

To investigate details of the latter Ca^2+^ signaling pathway, we first examined the contribution of LTCC-mediated transmembrane Ca^2+^ influx versus Ca^2+^ release from the ER to Ca^2+^ levels in either spontaneously firing hippocampal neurons and in neurons exposed to 10 Hz EFS, a stimulation paradigm leading to profound rises in mitochondrial Ca^2+^ (Dhoundiyal et al., 2022).

The inhibition of LTCC-mediated Ca^2+^ influx by isradipine (Isra, 1 μM) resulted in a reduction of both, spontaneously arising and EFS-induced Ca^2+^ levels in neuronal cell bodies. In the presence of isradipine, basal Ca^2+^ levels in spontaneously active neurons decreased by 28.99% (from 56.4 (±4.1) nM to 40.0 (±3.8) nM) and maximal (peak) cytosolic Ca^2+^ measured during 10 Hz EFS decreased by 60.68% (from 319.87 (±39) nM to 125.78 (±19.42) nM) (Fig. 1A, B).

**Figure 1:**
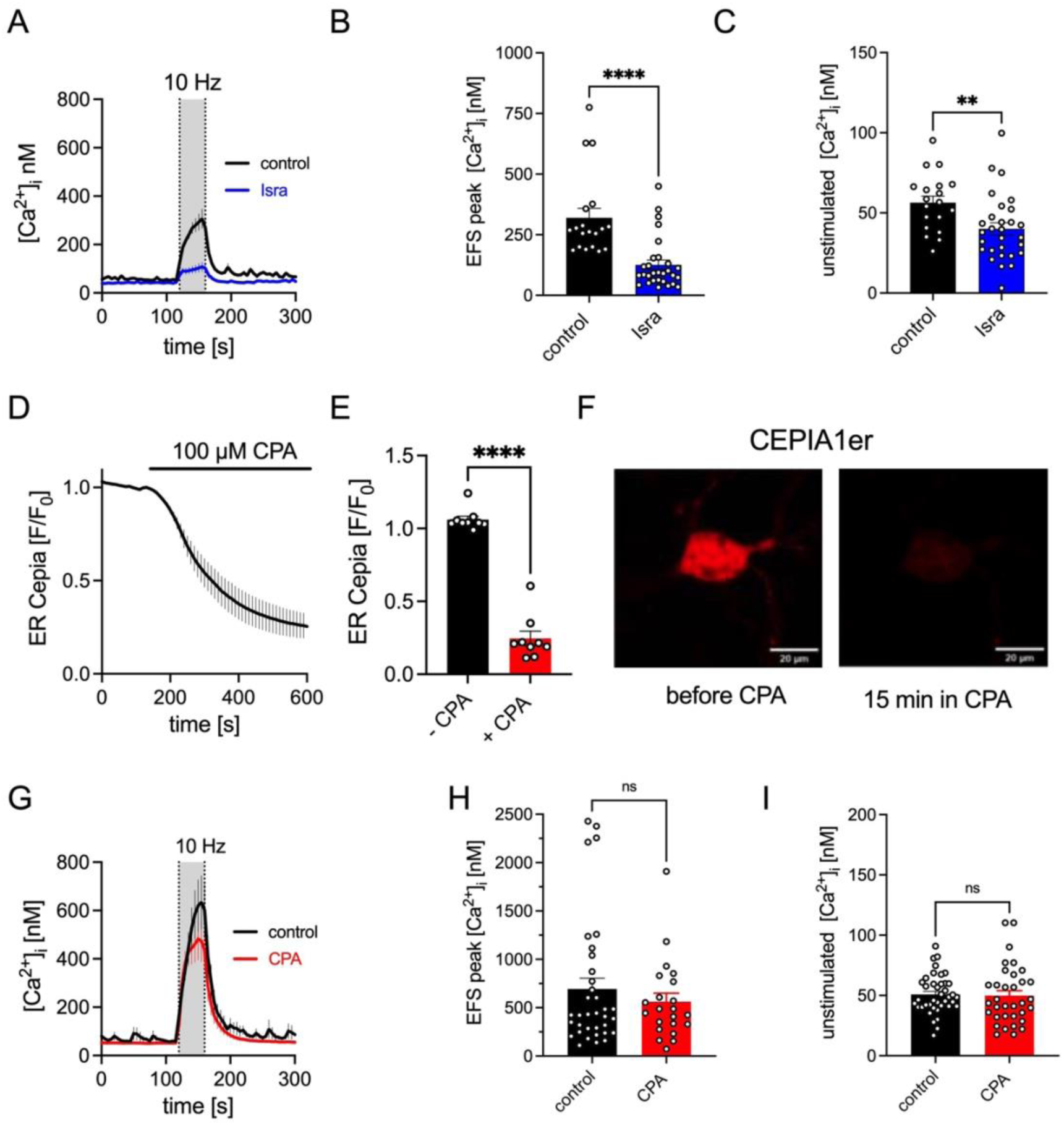
Contribution of extra– and intracellular Ca^2+^ stores to cytosolic Ca^2+^. (**A**) Changes in [Ca^2+^]_i_ measured using the Ca^2+^-sensitive dye Fluo4 AM. Neurons were electrically stimulated (EFS: 1.5 ms, 20 V for 40 s at 10 Hz, as indicated) in the presence (blue trace n=29 cells) or absence (black trace n=19 cells) of 1 μM isradipine. The fluorescent values obtained from individual neuronal somata were converted into [Ca^2+^]_i_, as described in the methods section. **(B,C)** Statistical evaluation of EFS-induced peak [Ca^2+^]_i_ values (B) and pre-stimulation [Ca^2+^]_i_ levels (C) extracted from the traces in panel A. **(D)** ER Ca^2+^ levels were monitored using the fluorescent ER Ca^2+^ sensor CEPIA1er. Mean fluorescent responses from 9 cells recorded upon application of 100 μM CPA. **(E)** Statistical evaluation of normalized CEPIA1er fluorescence before (black) and after (red) the application of CPA (n=9 cells from 2 preparations). **(F)** Representative micrographs of a neuron expressing CEPIA1er obtained before and after the application of CPA; Scale bar: 20 μm. **G)** Changes in [Ca^2+^]_I_ measured using the Ca^2+^-sensitive dye Fluo4 AM. Neurons stimulated by the EFS in the presence (red trace, n=22 cells) or absence (black trace, n=35 cells) of 100 μM CPA. **(H,I)** Statistical evaluation of maximal EFS-induced [Ca^2+^]_i_ values (H) and basal pre-stimulation [Ca^2+^]_i_ values (I) extracted from the traces in panel G. Data in panels B and H were evaluated using Mann–Whitney test, C (t(46)=2.836) and I(t(72)=0.1978) were compared with unpaired t-tests, and data in panel E (t(8)=14.13) were compared using paired t-test. [Ca^2+^]_i_: Cytosolic Ca^2+^ concentration; CPA: cyclopiazonic acid. **p < 0.01, ****p <0.0001.

To monitor the contribution of ER Ca^2+^ release, we used the specific SERCA inhibitor cyclopiazonic acid (CPA, 100 μM). CPA led to a gradual decrease in ER Ca²⁺ levels, as evidenced by reduced fluorescence of the ER Ca^2+^ indicator CEPIA1er. (Fig. 1D–F). ER Ca^2+^ levels reached a steady state after 8 minutes of CPA application indicating ER Ca^2+^ depletion. Nonetheless, ER Ca^2+^ depletion exhibited no discernible impact on baseline cytosolic Ca^2+^ concentrations (a reduction by 1.82% from 50.85 (±2.5) nM to 49.92 (±4.1) nM) (Fig. 1G, I). Likewise, ER Ca^2+^ depletion exerted only a marginal effect on the EFS-induced increase in cytosolic Ca^2+^. Following EFS, the peak cytosolic Ca^2+^ levels showed a slight, statistically insignificant decrease of 18.87% (from 693.18 (±112.6) nM to 562.30 (±87.2) nM) (Fig. 1H). These results suggest that activity-dependent elevations of cytosolic Ca^2+^ result preferentially from influx of extracellular Ca^2+^.

To further analyze a role of ER Ca^2+^ release, the two major ER Ca^2+^ release channels, RyRs and IP_3_Rs, were targeted by selective pharmacological tools. In hippocampal neurons in cell culture, RyRs and IP_3_Rs can be activated separately by caffeine and carbachol, respectively (Y. Wang et al., 2002). The rise in cytosolic Ca^2+^ as reflected by the increase in Fluo 4 fluorescence triggered by the muscarinic receptor agonist carbachol (CCh, 10 μM) was reduced when IP_3_Rs were blocked by either 30 μM or 100 μM of 2-aminoethoxydiphenyl borate (2-APB). Both concentrations of 2-APB were able to significantly dampen down the CCh-evoked increase in Fluo 4 fluorescence (Fig. 2A, B); however, the higher 2-APB concentration (100 μM) inhibited spontaneous fluctuations in the fluorescence signal as well, and the latter is correlated with spontaneous neuronal activity. As such an effect was not seen with partial IP_3_R inhibition by 30 μM 2-APB (Fig 2A, red trace), we used this lower 2-APB concentration for subsequent experiments. The caffeine-induced rise in cytosolic Ca^2+^ levels (Fig. 2C, D) was blocked completely by ryanodine (50 μM). Cytosolic Ca^2+^ levels determined before and during 10 Hz EFS remained unaffected by 50 μM ryanodine and were slightly increased by 30 μM 2-APB (Fig. 2E–G).

**Figure 2:**
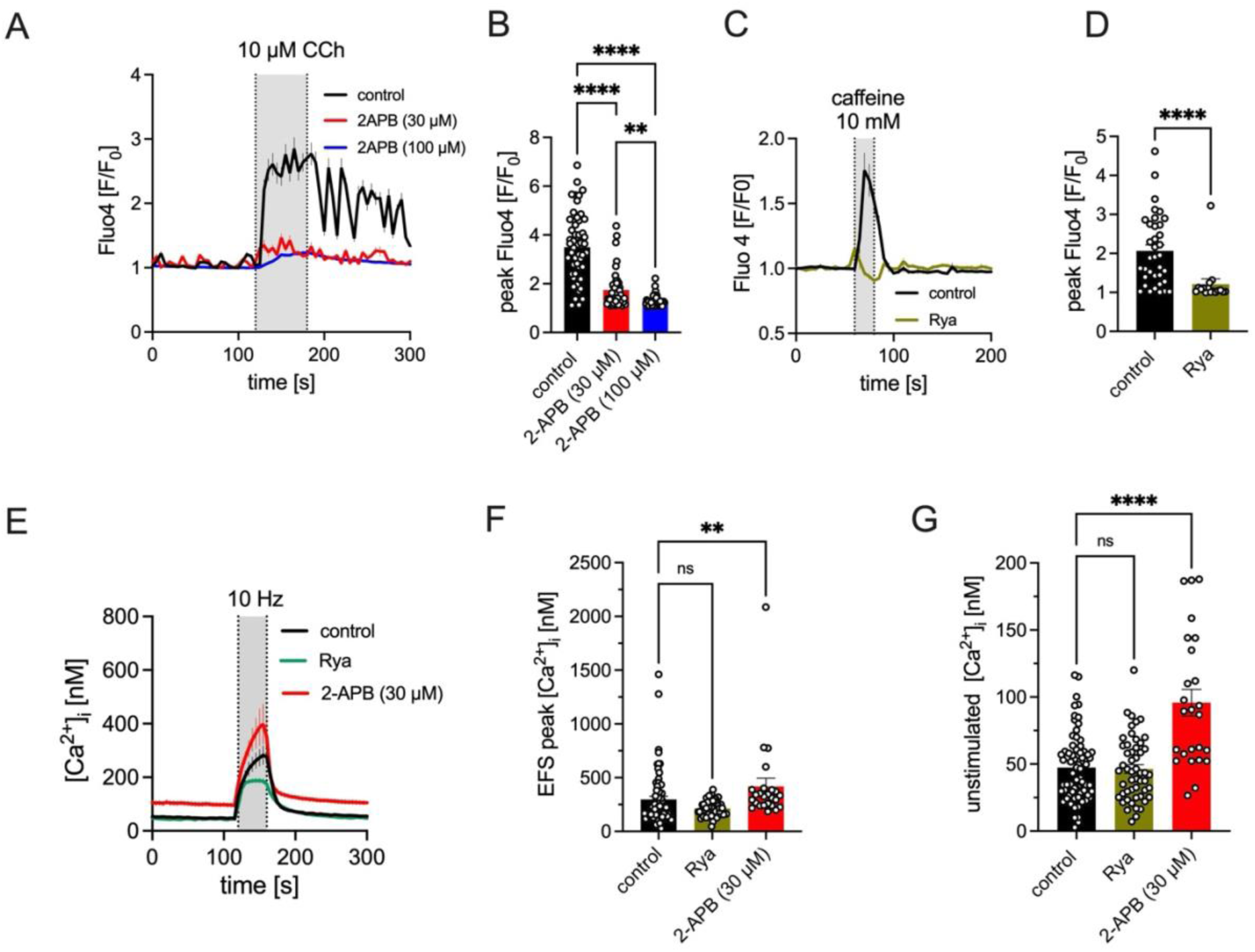
Effect of ER Ca^2+^ release channel inhibition on cytosolic Ca^2+^ dynamics. Changes in cytosolic Ca^2+^ levels obtained from neuronal somata were monitored in time using Fluo4 AM. **(A)** Averaged traces of cytosolic Ca^2+^ responses from neurons stimulated with 10 μM CCh in control buffer (black trace) or in the presence of 30 or 100 μM 2-APB (red and blue traces, respectively). **(B)** Statistical evaluation of the peak Fluo4 signals upon CCh stimulation from panel A. **(C)** Cytosolic Ca^2+^ response from neurons stimulated with 10 mM caffeine either in control buffer (black trace) or in the presence of 50 μM ryanodine (green trace). **(D)** Statistical evaluation of the peak Fluo4 signals upon caffeine stimulation shown in panel C. **(E)** Changes in [Ca^2+^]_i_ of EFS-stimulated neurons (EFS: 1.5 ms pulse of 20 V for 40 s at 10 Hz, as indicated) in control buffer (black trace) and in the presence of 50 μM ryanodine (green trace) or 30 μM 2-APB (red trace). The fluorescent values were converted into [Ca^2+^]_i_, as described in the methods section. **(F,G)** Statistical evaluation of maximal [Ca^2+^]_i_ values post-(F) and pre-(G) EFS stimulation of neurons shown in panel E. Data in panel B, F and G were analyzed using a Kruskal-Wallis one-way ANOVA for multiple comparisons. Data in panel D were statistically compared with Mann–Whitney test; [Ca^2+^]_i_: Cytosolic calcium concentration; 2-APB: 2-aminoethoxydiphenyl borate; CCh: Carbachol. **p < 0.01 and ****p <0.0001.

Altogether, these results indicate that under control conditions, Ca^2+^ enters the cytosol during neuronal activity preferentially from the extracellular space, and LTCCs represent a prominent Ca^2+^-entry pathway. ER-mediated calcium release provides only a minor contribution, if at all, to both, resting [Ca^2+^]_i_ levels as well as EFS-induced rises in cytosolic Ca^2+^.

### 3.2 ER Ca^2+^ depletion affects mitochondrial Ca^2+^ levels through IP_3_R-mediated Ca^2+^ release

Ca^2+^ coupling between ER and mitochondria may occur independently of changes in average cytosolic Ca^2+^ by relying on microdomains of high Ca^2+^(Rizzuto et al., 1993). At the MAMs, ER Ca^2+^ release channels are positioned near the MCU, where tethering proteins maintain their close association and facilitate the transfer of Ca^2+^ from the ER to the mitochondria. (Csordás et al., 2018; Hirabayashi et al., 2017; Rizzuto et al., 1993; Szabadkai et al., 2006).

Therefore, we monitored the direct impact of ER Ca^2+^ depletion on mitochondrial Ca^2+^ levels using the mitochondrially targeted Ca^2+^ sensor CEPIA2mt even though we had not found evidence for a contribution of ER-mediated Ca^2+^ release to the EFS-induced rise in cytosolic calcium. Depletion of the ER Ca^2+^ content by CPA decreased [Ca^2+^]_mito_ in both spontaneously firing and EFS-stimulated neurons (Fig. 3A-C). Basal mitochondrial calcium was reduced by 25.85 % (from 49.99 (±8.21) nM to 37.06 (±2.46) nM; Fig. 3C) and the maximal mitochondrial Ca^2+^ response to EFS by 51.19% (from 477.61(±126.1) nM to 233.10 (±30.82) nM; Fig. 3B). These findings indicate that transfer of Ca^2+^ from the ER to mitochondria may take place in neurons across a broad range of firing frequencies. Furthermore, most of the Ca^2+^ released from the ER during neuronal activity seems to be directed to mitochondria rather than the cytosol.

**Figure 3:**
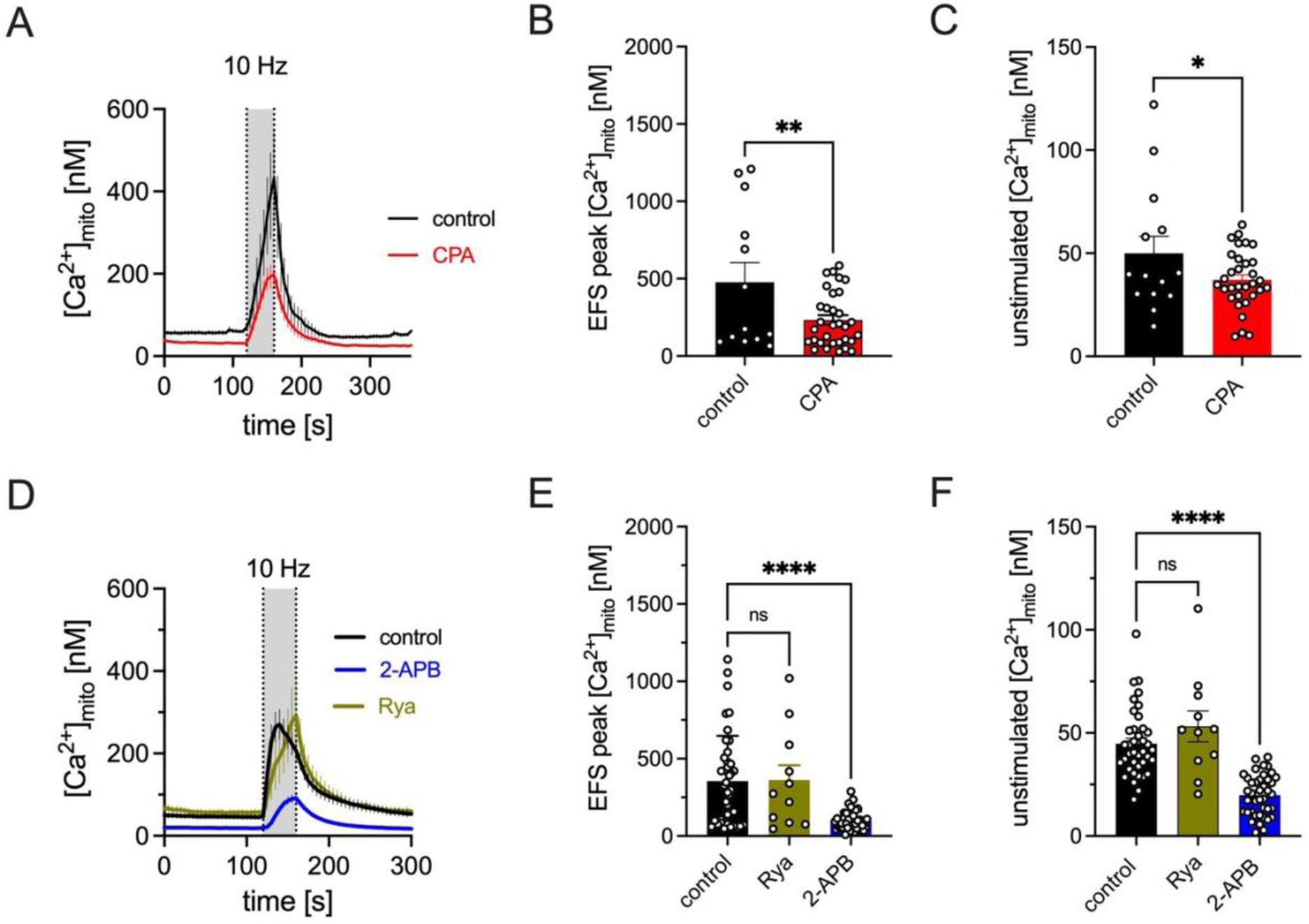
Contribution of ER Ca^2+^ release via IP_3_Rs to mitochondrial Ca^2+^ levels. Changes in [Ca^2+^]_mito_ were monitored in neurons expressing the fluorescent mitochondrial Ca^2+^ sensor CEPIA2mt. EFS of 10 HZ for 40 s was used to stimulate neuronal activity, each pulse being 1.5 ms and 20 V. The fluorescent values were converted into [Ca^2+^]_mito_ as described in the methods section. **(A)** Averaged [Ca^2+^]_mito_ response from the somata of hippocampal neurons upon EFS in the presence (red trace, n=33 cells) or absence (black trace, n=13 cells) of 100 μM CPA. **(B,C)** Peak [Ca^2+^]_mito_ values post stimulation (B), and basal pre-stimulation [Ca^2+^]_mito_ values (C) extracted from the data depicted in A. **(D)** Averaged [Ca^2+^]_mito_ response from the somata of hippocampal neurons upon EFS in control buffer (black trace) and in the presence of 50 μM Rya (green trace) or 30 μM 2-APB (blue trace), respectively. **(E,F)** Peak [Ca^2+^]_mito_ post stimulation values (E), and basal [Ca^2+^]_mito_ pre-stimulation values (F) extracted from the data depicted in D. Data in panel B (t(43)=2.635) and C (t(44)=1.976) were analyzed using unpaired t-test. Data in panel E were analyzed by Kruskal-Wallis one-way ANOVA and data in F (F(2,87)=35.43) were analyzed by one-way ANOVA followed by Dunnett’s test for multiple comparisons. [Ca^2+^]_mito_: Mitochondrial Ca^2+^ concentration; 2-APB: 2-aminoethoxydiphenyl borate; CPA: cyclopiazonic acid; *p < 0.05; **p < 0.01 and ****p <0.0001.

Both, RyRs and IP_3_Rs, have been shown to be localized at MAM sites, and both can be activated by Ca^2+^ (Hajnóczky et al., 2000, 2002). Therefore, both groups of channels have the potential to mediate Ca^2+^ transfer from the ER to the mitochondria. Inhibition of RyRs by ryanodine (50 μM) did not reduce basal mitochondrial Ca^2+^ levels (reduced by 6.8 %) (Fig. 3D, F), nor did it affect the EFS-induced mitochondrial Ca^2+^ transients (Fig. 3D green trace, 3E). Inhibition of IP_3_Rs by 30 μM 2-APB, in contrast, mimicked the effects of ER Ca^2+^ depletion by CPA on mitochondrial Ca^2+^ dynamics (Fig. 3d: blue trace). 2-APB decreased basal mitochondrial Ca^2+^ levels by 51.55 % (from 56.94 (±5.77) nM to 27.58 (±1.5) nM) (Fig. 3F) and peak mitochondrial Ca^2+^ transients by 56.27% (from 213.50 (±40.49) nM to 93.35 (±17.74) nM) (Fig. 3E). Therefore, it is IP_3_Rs rather than RyRs that mediate ER– mitochondria Ca^2+^ transfer.

### 3.3 IP_3_R-mediated ER–mitochondria Ca^2+^ transfer can contribute to ATP production in response to neuronal activity

The findings above suggest that ER–mitochondria Ca^2+^ transfer in hippocampal neurons is mediated by IP_3_Rs whether neurons are exposed to EFS or not. This leads to the question whether the transfer of Ca^2+^ from ER to mitochondria might be relevant for mitochondrial ATP production. To address this issue, we tracked ATP/ADP recovery following EFS of neurons expressing the genetically encoded ATP/ADP sensor PercevalHR, while ER Ca²⁺ release was modulated by pharmacological means.

Activity-dependent ATP production in neurons involves several competing metabolic pathways which are recruited according to the intensity of neuronal firing (Dhoundiyal et al., 2022). Furthermore, when one metabolic pathway is unavailable, others step in to compensate for this loss, and this ensures continuous ATP production and provides neurons with metabolic flexibility. In line with this idea, ER Ca^2+^ depletion by CPA had only a minor effect on the ATP/ADP recovery following EFS (Fig. 4A-C). Compared to the control neurons, the recovery rate (Fig. 4B) and the baseline Perceval fluorescence (Fig. 4C) in CPA-treated neurons decreased only slightly, presumably due to compensatory activation of Ca^2+^ sensitive redox shuttles (see below). Thus, the presence of multiple Ca^2+^-sensitive mechanisms makes studying the details of ER-mitochondria Ca^2+^ crosstalk rather challenging.

**Figure 4:**
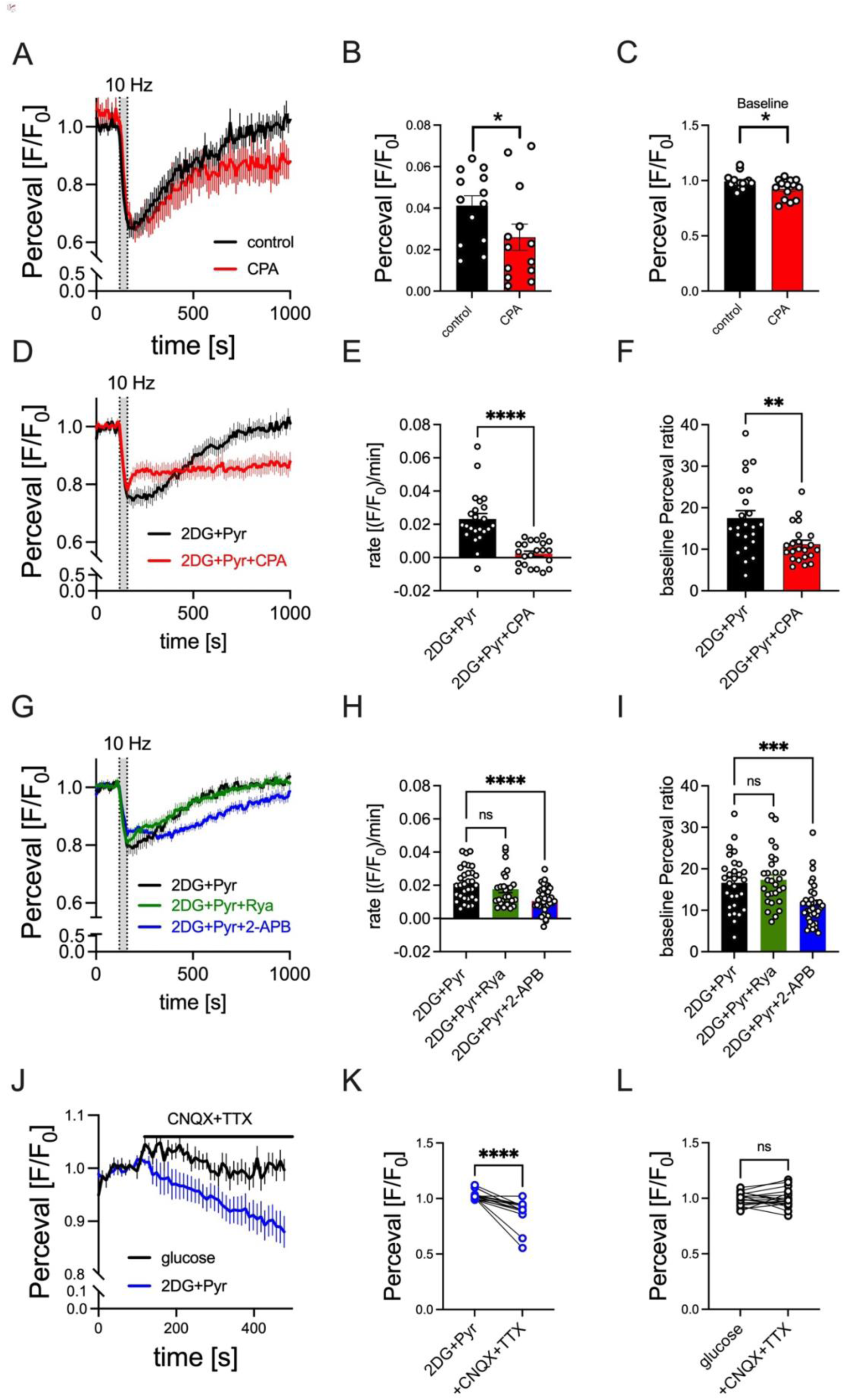
Contribution of ER Ca^2+^ release via IP_3_Rs to mitochondrial ATP synthesis. Cytosolic ATP/ADP was evaluated as the mean fluorescence of Perceval HR measured from neuronal somata. **(A-C)** Averaged traces (A) depicting the effect of CPA on EFS-induced rate of recovery of cytosolic ATP/ADP **(B)** as well as on PercevalHR baseline **(C)** n=13 cells. To isolate the role of mitochondrial calcium in stimulation of ATP synthesis, neurons were maintained in presence of 2 mM 2DG and 2 mM pyruvate. **(D)** Compared to control neurons (black trace), depletion of ER calcium by 100 μM CPA led to a marked decrease of the ATP recovery rate following EFS (red trace). **(E, F)** Statistical analysis of recovery rates (E) and basal, pre-stimulation Perceval ratios (F) n=22-23 cells. **(G-I)** Effect of RyR (green) or IP_3_R (blue) inhibition on the rate of ATP synthesis (H) and baseline Perceval ratios (I) n=26-37 cells. **(J-L)** Effect of inhibition of neuronal activity on cytosolic ATP/ADP ratio in neurons incubated in presence of 2DG+Pyr (J and K, blue, n-15 cells) and in control neurons incubated in external buffer (J and L, black, n-16 cells) Data in panels B (t(12)=2.316) and C(t(26)=2.326) were analyzed using paired t-test and data in panels E and F by Mann–Whitney test. Data in panel H (F(2,92)=9.809) were analyzed by one-way ANOVA followed by Dunnett’s test for multiple comparisons and data in panel I were analyzed by Kruskal-Wallis one-way ANOVA. Data in panels K and L were analyzed using paired Wilcoxon test. 2-APB: 2-aminoethoxydiphenyl borate; 2DG: 2-deoxyglucose; CPA: cyclopiazonic acid; Pyr: Pyruvate; *p<0.05,**p < 0.01; and ***p <0.001; ****p < 0.0001.

To overcome this problem, we aimed to experimentally enhance the bioenergetic dependence on ER-derived Ca²⁺ to enable a direct examination of the impact of ER Ca²⁺ on mitochondrial ATP production. To achieve this, the following experiments were performed in a glucose-free external buffer containing only the non-metabolizable glucose analogue 2-deoxyglucose (2DG, 2 mM) and the mitochondrial substrate pyruvate (pyr, 2 mM). Under these conditions, the redox shuttles regulated by extramitochondrial calcium (i.e. the malate aspartate shuttle and the glycerol 3-phosphate shuttle) cannot contribute to the mitochondrial ATP production and force mitochondria to rely on the stimulation of TCA cycle enzymes by matrix Ca^2+^. Additionally, this intervention circumvents the confounding contribution of ATP production by glycolysis (see Dhoundiyal et al., 2022).

In the presence of 2DG, neurons utilizing pyruvate fully recover their ATP/ADP ratios upon EFS (Fig. 4D, black trace), while depletion of the ER Ca^2+^ by CPA significantly slowed the rate of this ATP/ADP recovery (Fig. 4e, red trace). As a consequence, the majority of neurons were unable to restore their ATP/ADP ratio to pre-stimulation levels (Fig. 4D: blue trace). Perceval baseline fluorescence, reflecting the ATP/ADP levels in spontaneously active neurons, was also significantly lower in neurons incubated in 2DG+pyr+CPA as opposed to those incubated in 2DG+pyr alone (Fig. 4F).

To evaluate a potential impact of IP_3_R on mitochondrial ATP synthesis, neurons were exposed to EFS in 30 μM 2-APB plus 2DG+pyruvate: in the presence as compared to the absence of 2-APB, the rate of ATP recovery following EFS was diminished, which is consistent with decreased mitochondrial Ca^2+^ levels upon IP_3_R inhibition by 2-APB (Fig. 4G: blue trace, 4H). Moreover, 2-APB led to a decrease in baseline Perceval fluorescence ratios suggesting lower ATP/ADP levels in spontaneously firing neurons (Fig. 4I). In contrast, treatment of the neurons with ryanodine to inhibit RyRs had no influence on baseline ATP/ADP levels (Fig. 4I) nor did it alter ATP/ADP recovery after EFS (Fig. 4G-H: green).

The observation that disruption of ER-mitochondria Ca^2+^ crosstalk led to reduced ATP/ADP ratios (Fig. 4F, I) and lower mitochondrial calcium levels (Fig. 3C, F) even in absence of EFS suggests that this Ca^2+^ transfer may occur across a wide range of neuronal firing intensities. To investigate whether neurons incubated with 2DG and pyruvate can sustain their ATP/ADP balance through an activity-dependent stimulation of ATP synthesis even during spontaneous electrical activity (which corresponds to firing frequencies of about 0.3 Hz (Dhoundiyal et al., 2022)), we monitored ATP/ADP dynamics in spontaneously firing neurons. A combination of 10 μM CNQX and 0.5 μM TTX was applied to silence neuronal firing. When neurons were exposed to CNQX+TTX in 2DG+pyruvate incubated neurons, a gradual, continuous decline in Perceval fluorescence was observed (Fig. 4J, K, blue) which reflects a decrease in ATP/ADP ratio. Despite the removal of the energetic load imposed by neuronal firing, silent neurons exhibited a lower ATP/ADP ratio compared to their active counterparts. This suggests that neuronal activity promotes ER–mitochondria Ca^2+^ signaling, which is essential to adjust mitochondrial ATP production to energetic demands.

Consistent with findings from ER calcium depletion experiments, the dependence of mitochondrial ATP synthesis on neuronal activity was most evident when mitochondria relied on matrix calcium to stimulate metabolism (Figure 4J, K, blue). In conditions where multiple calcium-sensitive metabolic pathways were involved, ATP/ADP remained stable following the application of CNQX+TTX (Figure 4J and L, black).

### 3.4 Inhibition of PLC can decrease EFS-induced mitochondrial Ca^2+^ levels as well as the rate of ATP production

Although 2-APB is widely used in studying the involvement of IP_3_Rs in various cellular processes (Fernandes et al., 2022; Huang et al., 2017; Taha et al., 2024), the results obtained with this agent must be interpreted with caution. Since its introduction in 1997 as a potent IP_3_R blocker (Maruyama et al., 1997), the repertoire of its targets continues to expand. It has been shown to affect store-operated Ca^2+^ entry (Bilmen & Michelangeli, 2002; Bootman et al., 2002), SERCA pumps (Bilmen et al., 2002; Peppiatt et al., 2003), and transient receptor potential (TRP) channels (Hu et al., 2004; Ma et al., 2000), in addition to its effect on IP_3_Rs. Moreover, it has also been shown to interfere with functions of voltage-gated Ca^2+^ channels (Peppiatt et al., 2003). Similar problems come along with the use of xestospongin C, another widely used pharmacological IP_3_R inhibitor (Gambardella et al., 2021; Solovyova et al., 2002).

In order to corroborate a role of IP_3_Rs by independent means, inhibition of phospholipase C (PLC) was chosen as alternative method. This enzyme cleaves phosphatidylinositol 4, 5-bisphosphate (PIP_2_) into diacylglycerol (DAG) and IP_3_ and the latter then activates IP_3_Rs (Gomperts et al., 2009; Putney & Tomita, 2012). In cultures of hippocampal neurons, the cytosolic rise of Ca^2+^ in response to EFS relies on the release of endogenous glutamate (Dhoundiyal et al, 2022). Hence, released glutamate can be expected to stimulate PLC through activation of mGluRs.

To support this conjecture, we monitored PIP_2_ cleavage which is the substrate for PLC to synthetize IP_3_. To this end, a BFP-tagged pleckstrin homology domain of PLCδ that specifically binds to PIP_2_ as a reporter (BFP-PH, see methods) (Fig. 5A-C) was expressed. Under baseline conditions, this reporter is predominantly localized to the neuronal plasma membrane where PIP_2_ resides. Upon activation of PLC, PIP_2_ is lost from the membrane due to enzymatic cleavage, and the PH domain is released into the cytosol which leads to a shift in fluorescence. To accurately measure the decrease in BFP-PH fluorescence resulting from PIP_2_ cleavage—and to correct for potential fluorophore bleaching—we quantified the ratio of membrane-associated to cytosolic fluorescence signals. Upon EFS, we observed a rapid decrease of the reporter fluorescence ratio with some partial post stimulation recovery (Fig. 5B, C). This result demonstrates that 10 Hz EFS leads to PIP_2_ cleavage most likely through a receptor-dependent activation of PLC which generates IP_3_.

**Figure 5:**
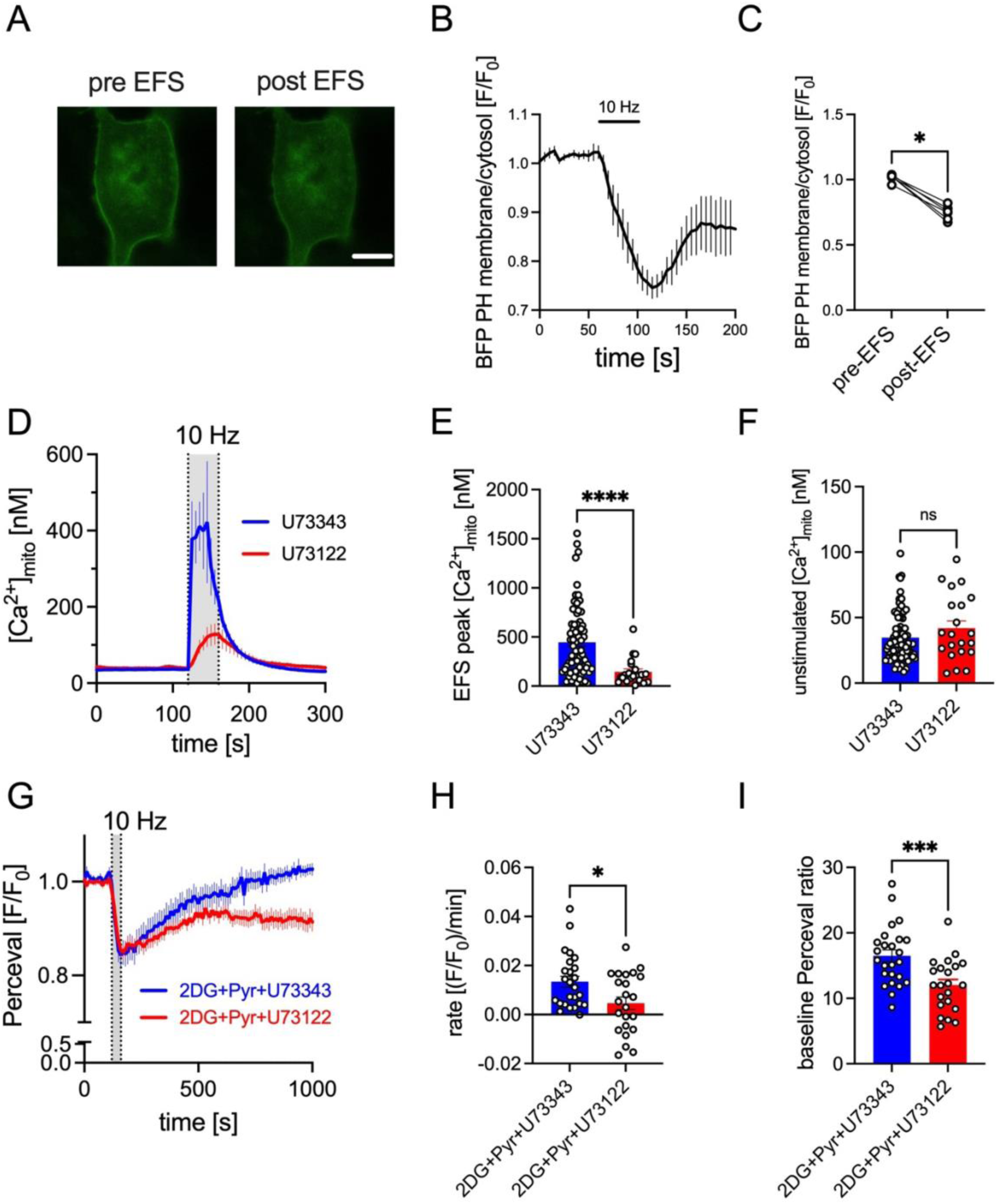
Effect of the inhibition of PLC-mediated PIP_2_ cleavage on mitochondrial Ca^2+^ levels and ATP production. Plasma membrane PIP_2_ levels were monitored in hippocampal neurons expressing BFP-PH reporter. Changes in [Ca^2+^]_mito_ were monitored in neurons expressing the fluorescent mitochondrial Ca^2+^ sensor CEPIA2mt. EFS of 10 Hz for 40 s was used to stimulate neuronal activity, each pulse being 1.5 ms and 20 V. Changes in the CEPIA2mt fluorescence upon EFS were recorded. The fluorescent values were converted into [Ca^2+^]_mito_ as described in the methods section. (**A**) Original micrographs illustrating BFP-PH signal pre– and post-EFS. Scale bar = 7 μm. (**B-C**) Monitoring of effect of EFS on a ratio of membrane and cytosolic BFP-PH fluorescence (B) with its statistical analysis (C) (n = 7 cells). **(D)** Averaged [Ca^2+^]_mito_ response from somata of hippocampal neurons upon EFS in the presence of U73122 (red trace) or U73343 (blue trace) (preincubated for 1 h). **(E, F)** Peak [Ca^2+^]_mito_ values post stimulation (E), and basal [Ca^2+^]_mito_ pre-stimulation values (F) extracted from the traces depicted in D. **(G)** Time course of normalized Perceval values obtained in neurons incubated in 2 mM 2DG+2 mM pyruvate+1 μM U73343 ( blue trace, n=28 cells), or 2DG+pyruvate+1 μM U73122 (red trace, n=30 cells). **(H)** Rate of recovery of the fluorescent values after stimulation (calculated as described in the methods section) were compared between the two conditions shown in (G). **(I)** Comparison of the baseline Perceval fluorescence values derived from neurons in (G) prior to stimulation. Data on panel C were statistically compared using Wilcoxon test. Data on panel E, F, H and I were compared with Mann–Whitney test. [Ca^2+^]_mito_: mitochondrial calcium concentration; 2DG: 2-deoxyglucose; Pyr: Pyruvate; PIP_2_: phosphatidylinositol 4, 5-bisphosphate *p < 0.05; ***p < 0.001; ****p < 0.0001.

To corroborate this latter mechanism, neurons were exposed to the PLC inhibitor U73122 or its inactive analogue U73343 before, during, and after EFS. This approach has been shown to prevent Ca^2+^ release from intracellular stores in a variety of cell types (Jin et al., 1994; Tatrai et al., 1994). Compared to neurons incubated in U73343, those exposed to U73122 showed reduced mitochondrial calcium uptake following EFS (Fig. 5D, E). Basal mitochondrial calcium levels, in contrast, were unaffected (Fig. 5F). In line with the reduction in mitochondrial Ca^2+^, neurons incubated in U73122 showed a slower rate of ATP/ADP ratio recovery as compared to those in U73343 (Fig. 5G: blue trace, 5H). As a consequence, upon EFS, neurons treated with U73122 failed to restore their ATP levels to those seen before stimulation (Fig. 5G). The baseline ATP levels were also considerably lower in the presence of U73122 as compared with U73343 (Fig. 5I). These results confirm that IP_3_R-mediated ER Ca^2+^ release stimulates mitochondrial ATP synthesis to support neuronal metabolism during electrical activity.

## 4. Discussion

The essential role of Ca^2+^ transfer from the ER to mitochondria in maintaining basal cellular bioenergetics has been demonstrated across various cell types. In most cells, this Ca^2+^ transfer is mediated by close interactions of IP_3_Rs and mitochondrial voltage dependent anion carriers (VDACs) located at the outer mitochondrial membrane (Bartok et al., 2019; Cárdenas et al., 2010, 2016; Cruz et al., 2021; Suman et al., 2018). For this reason, pharmacological or genetic suppression of IP_3_R activity interferes with both, the transport of Ca^2+^ and the synthesis of ATP within mitochondria (Cardenas et al., 2020; Cruz et al., 2021).

However, ER–mitochondrial Ca^2+^ transfer and its involvement in mitochondrial bioenergetics in neurons have received relatively little attention. While existing studies underscore the importance of ER-mitochondria Ca^2+^ transfer in neuronal bioenergetics, there still is inconsistency with respect to the specific types of ER Ca^2+^-release channels that mediate this process. In a study on cortical neurons, RyRs, but not IP_3_Rs, were demonstrated to control basal cellular respiration. Notably, this effect did not result from direct ER-mitochondria Ca²⁺ transfer, as RyR-mediated stimulation of cellular respiration occurred independently of mitochondrial Ca²⁺ uptake (Pérez-Liébana et al., 2022). Another study investigated spontaneously spiking dopaminergic neurons in the substantia nigra and found that stimulation of mitochondrial ATP synthesis during spontaneous spiking was mediated by RYRs. However, the authors noted that in principle, both types of ER Ca²⁺ release channels could facilitate ER-mitochondria coupling (Zampese et al., 2022). In addition, a recent study on Wolfram syndrome, a rare genetic disease caused by mutations in the WFS1 or CISD2 gene, in primary cortical neurons found that an inefficient IP_3_R-mediated ER–mitochondria Ca^2+^ exchange due to reduced ER Ca^2+^ content could drive disease progression (Liiv et al., 2024). Discrepancies in these findings may arise from methodological variations and inherent differences between neuronal subtypes. In addition to these inconsistencies, most of these studies have evaluated the role of ER Ca^2+^ release in neuronal metabolism under relatively uncontrolled conditions. Key factors, such as the intensity of the underlying neuronal activity, have not been monitored, and stimulation protocols have employed rather non-physiological ways of neuronal activation (e.g. high concentrations of NMDA, caffeine).

Another factor complicating the interpretation of the results obtained is the fact that neurons display a high degree of metabolic flexibility, and inhibition of one metabolic pathway leads to a compensatory increase in the activity of other pathways. For instance, reducing MCU-mediated calcium uptake results in a compensatory rise in the activity of cytosolic redox shuttles, thereby enhancing their role in maintaining the overall bioenergetic balance (Zampese et al., 2022; Dhoundiyal et al., 2022). Moreover, engagement of these metabolic pathways is modulated by the intensity of neuronal firing (Dhoundiyal et al., 2022, (Groten & MacVicar, 2022; Stoler et al., 2022). As a result, spontaneous neuronal firing and the recruitment of various metabolic pathways may substantially differ from one study to another. All of these factors make investigations regarding the involvement of ER-mitochondria Ca^2+^ transfer in neurons complex. With the current approach, we did not aim to quantitatively assess which Ca^2+^ sensitive metabolic pathway might dominate in stimulating ATP synthesis during highly variable spontaneous neuronal firing; rather, our goal was to determine whether the calcium transfer from the ER to mitochondria can effectively stimulate ATP synthesis and if so, which ER Ca^2+^ release channel(s) might be involved.

To accomplish this, we reduced the likeliness of compensatory changes by selecting a pharmacological over a genetic approach, with all interventions in this study conducted acutely. In addition, key experiments were executed under conditions in which neurons relied solely on the stimulatory action of matrix Ca^2+^ on mitochondrial metabolism; by employing 2DG and pyruvate, we achieved reduced glycolytic flux, deprived redox shuttles from NADH, and maintained mitochondrial OXPHOS functional through pyruvate supplementation (Dhoundiyal et al., 2022).

Electric field stimulation as used in this study was selected as it offers good control over the intensity of induced neuronal firing. In addition, similar to endogenous neuronal activity, EFS relies on synaptic transmission and the propagation of action potentials, thereby mimicking the physiological way of neuronal activation (Dhoundiyal et al., 2022). The 10 Hz stimulus frequency was chosen based on our former experiments due to its capability of inducing significant mitochondrial calcium influx (Dhoundiyal et al., 2022).

The present results show that mitochondrial ATP production in neurons can be maintained through an activity-dependent, IP_3_-mediated transfer of Ca^2+^ from the ER to mitochondria within a wide range of neuronal firing frequencies. This was demonstrated by the actions of 2-APB, a purportedly selective inhibitor of IP_3_Rs (Maruyama et al., 1997). However, 2-APB was reported to modulate functions in a variety of ion channels including various TRP channels (Singh et al., 2018). TRPV channels, in particular, are activated by 2-APB (Singh et al., 2018), and TRPV1 (Hurtado-Zavala et al., 2017) as well as TRPV4 (Shibasaki et al., 2007) can mediate depolarizations in hippocampal neurons. This may explain why application of 2-APB led to increases in cytosolic Ca^2+^ concentrations which, however, were not accompanied by a rise, but rather by a decline, in mitochondrial Ca^2+^. Hence, activation of TRPV channels is unlikely to contribute to effects of 2-APB on mitochondrial Ca^2+^ and ATP production during neuronal firing. Moreover, respective actions of 2-APB were confirmed by an independent approach, namely inhibition of PLC and a resulting decline in IP_3_ synthesis. RyR blockage by ryanodine, in contrast, failed to affect both, ER-mitochondria Ca^2+^ transfer and ATP synthesis, whether neurons were exposed to EFS or not.

Due to the presence of multiple compensatory metabolic pathways, it is challenging to accurately determine the relative contribution of ER-mitochondria Ca²⁺ crosstalk to overall ATP synthesis. The findings in neurons not exposed to 10 Hz EFS in this study along with our previously published results on neurons exposed to 2.5 Hz EFS (Dhoundiyal et al., 2022) indicate that ER-mitochondria Ca²⁺ transfer may operate during low-intensity neuronal firing, even though redox shuttles are likely to play a more dominant role under these conditions. I addition, the data suggest that the importance of ER-mitochondria Ca²⁺ crosstalk increases with rising neuronal firing intensities (see figure 4E in Dhoundiyal et al., 2022) up to the point where cytosolic Ca²⁺ fluctuations become sufficiently large to bypass ER-mediated release and to directly trigger mitochondrial Ca²⁺ uptake (Hotka et al., 2020). Accordingly, neurons can utilize activity-driven ER-mitochondria Ca^2+^ transfer in wide range of firing intensities, but this mechanism is competing with other Ca^2+^ sensitive metabolic pathways.

### 4.1 Limitations

It should be noted that our results reflect mitochondrial ATP synthesis and associated processes in neuronal somata. However, as previous findings have indicated, these processes may vary in different subcellular compartments, such as dendritic spines, dendritic shafts, axonal segments, etc., due to the complexity of neuronal architecture (Ashrafi et al., 2020; Heine et al., 2020). Hence, the mechanisms by which Ca^2+^ regulates metabolism in neuronal compartments other than somata might differ from the results reported here. In addition, the coupling between neuronal activity and ER-mitochondria Ca^2+^ exchange may vary across different neuronal subtypes. An example of this is the spontaneously spiking dopaminergic neurons of the substantia nigra, which exhibit electrophysiological properties distinct from those of the hippocampal neurons examined in this study (Zampese et al., 2022). Studying these mechanisms in various neuronal subtypes is therefore essential. Another limitation is the use of 2DG + pyruvate to study the role of ER-mitochondria Ca^2+^ transfer in ATP synthesis. While these conditions are artificial, they allow us to isolate this metabolic pathway and thereby enable more robust conclusions about underlying mechanism.

## 5. Conclusion

In summary, the results obtained in hippocampal neuronal networks derived from new born rats, demonstrate an involvement of ER Ca^2+^ in activity-dependent mitochondrial metabolism. While LTCCs contribute little to the maintenance of basal cytosolic Ca^2+^ levels, their share in Ca^2+^ rise increases significantly upon EFS. Additionally, neurons can exploit a constitutive, activity-driven, Ca^2+^ flow from intracellular Ca^2+^ stores into mitochondria. During neural stimulation and ongoing CICR, this delivery of Ca^2+^ into the mitochondrial matrix boosts ATP production. In somata of rat hippocampal neurons, it is IP_3_Rs, but not RyRs, that mediate this transfer. Obviously, this may not apply to other neurons and subcellular neuronal compartments. Thus, alternative mechanisms of ER to mitochondrial Ca^2+^ transfer remain open for future investigations.

## CRediT authorship contribution statement

**Ankit Dhoundiyal**: Investigation, Formal analysis, Data Curation, Writing – original draft.

**Vanessa Goeschl**: Methodology, Writing – review and editing

**Stefan Boehm**: Writing – review and editing

**Helmut Kubista** – Methodology, Writing – review and editing, Funding acquisition

**Matej Hotka** – Conceptualization, Investigation, Formal analysis, Writing – original draft, Project administration, Supervision, Funding acquisition

## Availability of data and material

The authors confirmed that the data supporting the current study are available within the article and supplementary material. The raw data are also available from the corresponding author, upon reasonable request.

