## Supplemental Figures S1 and S2 for "IP_3_-mediated Ca^2+^ transfer from ER to mitochondria stimulates ATP synthesis in primary hippocampal neurons"

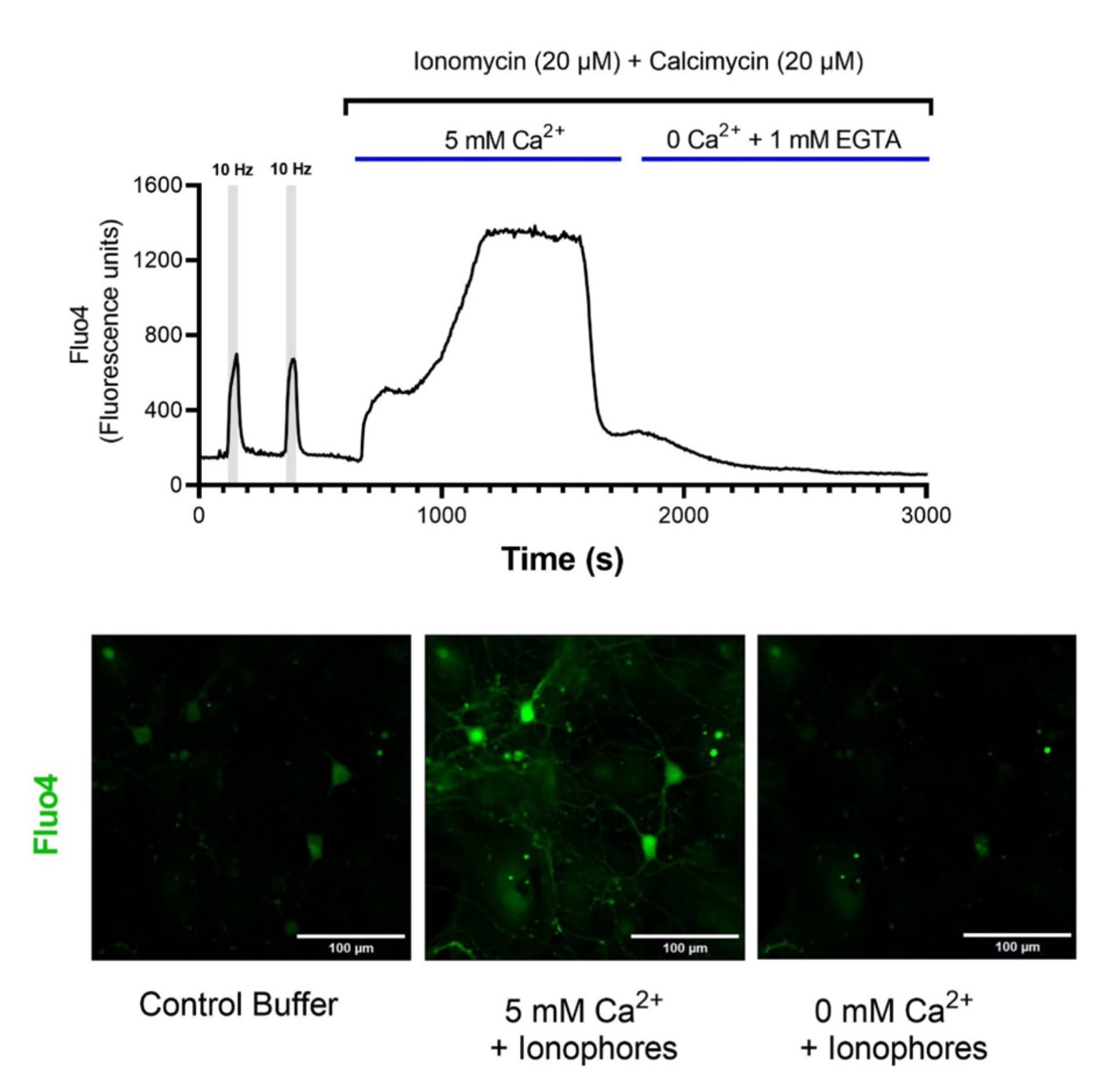

***Figure S1: Calculating F_max_ and F_min_ in neurons incubated in Fluo4 AM:*** ***a)*** *Time course of a representative Fluo4 response from a neuron subjected to two 10 Hz EFS, followed by the application of ionophores. Cells were flushed with a ionophore-containing solution with a high Ca^2+^ concentration (5 mM) in order to calculate F_max_. After that, the fluorescence was allowed to stabilize, which usually took 15 to 20 minutes. Following this, cells were washed with a 0 Ca^2+^ solution that contained ionophores and 1 mM EGTA. Once more, the fluorescence was given 15 to 20 minutes to stabilize (which gives F_min_) before the recording was stopped.* ***b)*** *Representative micrographs showing the Fluo-4 fluorescence in neurons incubated with control external buffer, high Ca^2+^ solution + ionophores and zero Ca^2+^ solution + ionophores.*

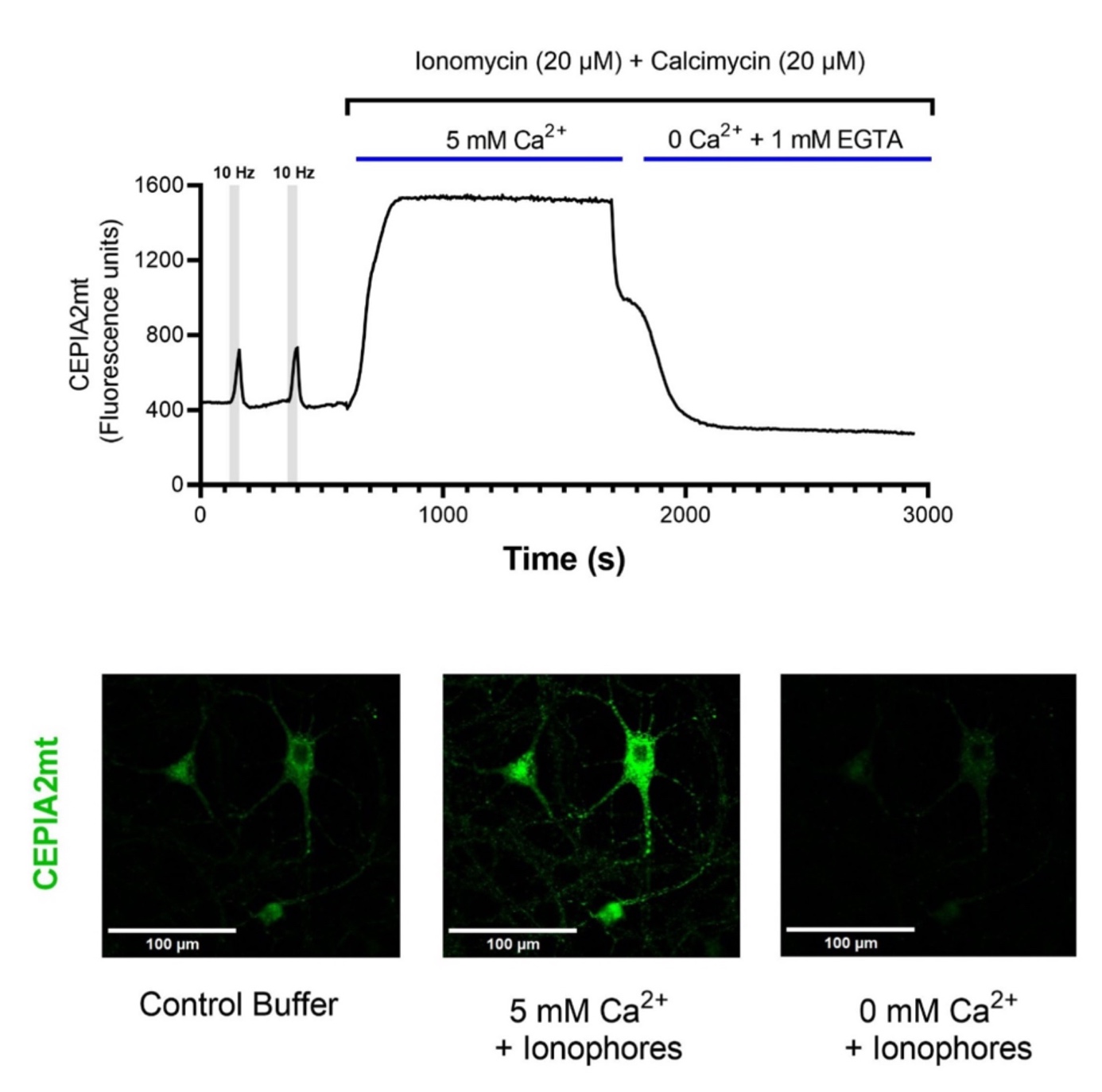

***Figure S2: Calculating F_max_ and F_min_ in a neuron expressing CEPIA2mt:*** ***a)*** *Time course of a sample CEPIA2mt response from a neuron subjected to two 10 Hz EFS, followed by the application of ionophores. In order to calculate F_max_, cells were flushed with a high Ca^2+^ concentration (5 mM) solution containing ionophores. The fluorescence was then allowed to reach a stable value, which took typically around 15-20 minutes. After this, cells were flushed with 0 Ca^2+^ solution containing 1 mM EGTA and ionophores. The fluorescence was again allowed to stabilize for 15-20 minutes before ending the recording.* ***b)*** *Representative micrographs showing a neuron expressing CEPIA2mt incubated with control external buffer, high Ca^2+^ solution + ionophores and zero Ca^2+^ solution + ionophores.*
